# Basal forebrain cholinergic neurons selectively drive coordinated motor learning in mice

**DOI:** 10.1101/2021.04.24.441273

**Authors:** Yue Li, Edmund Hollis

## Abstract

Motor control requires precise temporal and spatial encoding across distinct motor centers that is refined through the repetition of learning. The coordination of circuit refinement across motor regions requires modulatory input to shape circuit activity. Here we identify a role for the basocortical cholinergic pathway in the acquisition of a coordinated motor skill in mice. Targeted depletion of basal forebrain cholinergic neurons results in significant impairments in training on the rotarod task of coordinated movement. Cholinergic neuromodulation is required during training sessions as chemogenetic inactivation of cholinergic neurons also impairs task acquisition. Rotarod learning drives coordinated refinement of corticostriatal neurons arising in both medial prefrontal cortex (mPFC) and motor cortex, and we have found that cholinergic input to both motor regions is required for task acquisition. Critically, the effects of cholinergic neuromodulation are restricted to the acquisition stage, as depletion of basal forebrain cholinergic neurons after learning does not affect task execution. Our results indicate a critical role for cholinergic neuromodulation of distant cortical motor centers during coordinated motor learning.

## Introduction

The learning of coordinated motor behavior requires synchronization across distinct motor centers throughout the central nervous system. Coordinated task acquisition drives dynamic patterns of neuronal activity and synaptic plasticity in the cortex and striatum. Within the striatum, there is a progressive shift in regional influence, with dorsomedial striatum critical in early stages, followed by a shift to dorsolateral striatum later in task acquisition (Yin et al., 2009). Coherence develops within corticostriatal motor networks with precise timing of primary motor cortex (M1) and dorsolateral striatum (DLS) activity in support of motor performance across learning (Koralek et al., 2013; Koralek et al., 2012; Lemke et al., 2019). During accelerating rotarod training, associative medial prefrontal cortex connections to dorsomedial striatum (mPFC-DMS) are recruited concurrently alongside motor circuits projecting from primary motor cortex to dorsolateral striatum (Kupferschmidt et al., 2017). Early in learning, mPFC-DMS and M1-DLS circuits show parallel activation, while these patterns diverge as animals master the motor skill (Kupferschmidt et al., 2017). In rodents that have mastered learned motor skills, there is a dissociation between M1 and movement execution. On simple tasks, M1 appears to be dispensable as complete ablation does not alter trained movements (Kawai et al., 2015); whereas, more complex motor tasks have demonstrated that M1 activity is required for the appropriate execution of fine motor control (Lemke et al., 2019). Modulated motor control arises during motor training with M1 and DLS contributing to gross movement, while only M1 inactivation disrupts fine movements (Lemke et al., 2019). Neuromodulation of M1, but not mPFC, via cholinergic input from basal forebrain has been shown to be a key mechanism that supports the cortical plasticity that occurs during skilled motor learning in the rat (Conner et al., 2010). We have assessed the role of cholinergic input to M1 and mPFC in coordinated motor control.

The basal forebrain is one of the principal sites of acetylcholine synthesis in the brain. Cholinergic projections arising from basal forebrain subregions innervate distinct targets, modulating functions varying from motor control, sensory and perceptual coding, attention, memory, to anxiety (Boskovic et al., 2019; Zaborszky et al., 2018). Cholinergic neurons in the nucleus basalis of Meynert (NBM) have widespread projections to the cerebral cortex, while neurons in the diagonal band of Broca (DBB) and substantia innominata (SI) send projections to the prefrontal cortex (Boskovic et al., 2019). Basal forebrain cholinergic neurons are consistently activated at the onset of movement (Harrison et al., 2016), potentially synchronizing distant cortical motor centers. Neuromodulation requires temporal precision to modulate target area dynamics and synaptic plasticity, and reinforce cognitive and reward behaviors (Gil et al., 1997; Kruglikov and Rudy, 2008; Metherate et al., 1992).

Cholinergic signaling throughout cortical sensory and motor regions acts to modulate network properties and enhance glutamatergic signaling, driving long-lasting changes in synaptic strength (Rasmusson, 2000). In the auditory cortex, muscarinic acetylcholine signaling plays an instructional role in receptive field plasticity (Bakin and Weinberger, 1996; Kilgard and Merzenich, 1998; Miasnikov et al., 2001), while in visual cortex, cholinergic innervation is required for experience-dependent visual plasticity during the critical period (Bear and Singer, 1986). Following critical period closure, visual cortex plasticity is limited in adulthood by the prototoxin lynx1, an allosteric regulator of nicotinic acetylcholine receptors that reduces acetylcholine sensitivity (Miwa et al., 2006). Knockout of *lynx1* extends ocular dominance plasticity beyond the critical period into adulthood (Morishita et al., 2010). Within M1, the maturation of motor representations, or maps, depends upon basal forebrain cholinergic input (Ramanathan et al., 2015). In rats, basal forebrain cholinergic input to M1 has been found to support the plasticity of these motor maps as well as the remodeling of corticospinal neuron dendrite morphology (Conner et al., 2003; Conner et al., 2010; Wang et al., 2011). Across sensory modalities, the basal forebrain cholinergic system acts as a rapid and precisely timed reinforcement signal to support fast cortical activation and plasticity in associative learning (Hangya et al., 2015; Hanson et al., 2021; Liu et al., 2015).

In the present study, we demonstrate a key role for cholinergic basal forebrain neurons in the acquisition of coordinated motor behavior. We used targeted toxin-mediated and genetic depletion of NBM/SI cholinergic neurons and found that the acquisition of accelerating rotarod behavior was impaired. Disrupting the temporal activation of NBM/SI cholinergic neurons similarly impaired task acquisition. As rotarod task acquisition requires coordinated activity across distant motor centers, we tested whether selectively targeting cholinergic innervation of mPFC or primary motor cortex affected performance and found that cholinergic input to both areas was required for coordinated motor learning. The effects of disrupting basal forebrain cholinergic input to the cortex was specific to the acquisition phase of rotarod behavior as we found that cholinergic depletion after training elicited no effect on task execution. In contrast to previous findings in the rat (Conner et al., 2003), we found no effect of cholinergic depletion on the learning of a skilled forelimb task. Our findings demonstrate a critical role for cholinergic neuromodulation in coordinated motor performance.

## Materials and Methods

### Mice

All animal experiments and procedures were approved by the Weill Cornell Medicine Institutional Animal Care and Use Committee. All mice were housed on a 12-hour light/dark cycle from 6am to 6pm at 25°C with free access to food and water. For skilled pellet reach behavior, animals were calorie restricted to 80–90% of their free-feeding bodyweight. Male and female C57BL/6 animals (8-12 weeks old) were purchased from Jackson Laboratory. ChAT-Cre mice were originally obtained from Jackson Laboratory and bred in-house. In our study, hemizygous ChAT-Cre mice were used, which were achieved by back-crossing the homozygous ChAT-Cre line to C57BL/6 mice. In some experiments, hemizygous *ChAT-Cre::Ai14* mice were used. This mouse line expresses tdTomato in ChAT-positive neurons.

### Cholinergic neuron ablation by p75-saporin

Twenty-eight ten-week-old C57BL/6J mice were anesthetized with 4% isoflurane, maintained during surgery with 1.5-3% isoflurane, and had body temperature maintained at 37°C using a SomnoSuite small animal anesthesia system (Kent Scientific). Subcutaneous (SQ) injection of buprenorphine (0.1 mg/kg) and meloxicam (2 mg/kg) was given immediately following anesthesia. For global ablation of cholinergic neurons in NBM/SI basal forebrain, anti-p75 conjugated saporin (p75-saporin) or IgG-saporin control (*n* = 8 / group, Advanced Targeting Systems) was diluted to final concentration of 0.4 mg/ml in normal saline. Saporin solution (150 nl/site) was bilaterally injected into NBM/SI areas at a rate of 120 nl/min using a glass micropipette filled with mineral oil and attached to a programmable Nanoject III (Drummond Scientific). NBM/SI injection sites: 1) A/P -0.22 mm, M/L ±1.25 mm, D/V -4.7 mm; 2) A/P -0.7 mm, M/L ±1.75 mm, and D/V -4.7 mm. Following each injection, the pipette was held in place for an additional 4 min to prevent backflow. Mice were allowed to recover from surgery for 2 weeks prior to behavioral experiments.

In order to selectively ablate basal forebrain cholinergic neurons projecting to specific cortical regions, focal injections were performed in the target areas. IgG-saporin or p75-saporin was diluted to a final concentration of 0.08 mg/ml in normal saline. Saporin solution was bilaterally injected into mPFC, motor cortex, or visual cortex at a rate of 40 nl/min. The following injection volume and coordinates of injection sites were used: mPFC site 1) 200 nl at A/P +2.3 mm, M/L ±0.3 mm, D/V –1.5 mm; mPFC site 2) 200 nl at A/P +1.5 mm, M/L ± 0.3 mm, D/V -2.0 mm; motor cortex site 1) 200 nl at A/P +1.5 mm, M/L ±1.3 mm, D/V -0.6 mm; motor cortex site 2) 200 nl at A/P +0.5 mm, M/L ±1.5 mm, D/V -0.6 mm; motor cortex site 3) 200 nl at A/P -0.5 mm, M/L ±1.2 mm, D/V -0.6 mm; visual cortex site 1) 250 nl at A/P -2.8 mm, M/L ±2.3 mm, D/V -0.6 mm; visual cortex site 2) 250 nl at A/P -3.8 mm, M/L ±2.3 mm, D/V -0.6 mm.

### Cholinergic neuron ablation by diphtheria toxin

A targeted genetic approach was used to ablate basal forebrain cholinergic neurons. Sixteen eight-week-old transgenic mice expressing Cre recombinase behind the ChAT promoter crossed with Ai14 *Rosa-LSL-tdTomato* in the C57BL/6J background (*ChAT-Cre::Ai14*) were transduced with AAV encoding Cre recombinase-dependent diphtheria toxin receptor and the fluorescent reporter eYFP (AAV-DJ/8-EF1A-FLEX -DTR-P2A-EYFP) by bilateral injection into NBM/SI at a rate of 120 nl/min (Herman et al., 2016). AAV expressing only Cre-dependent eYFP fluorescent protein (AAV-DJ/8-EF1A-FLEX-P2A-EYFP) was used as control. NBM/SI was injected with 350 nl of virus [8.73×10^12^ VG/ml] at the following coordinates: site 1) A/P -0.1 mm, M/L ±1.25 mm, D/V -4.7 mm; site 2) A/P -0.45mm, M/L ±1.5 mm, D/V -4.7 mm; site 3) A/P -0.8 mm, M/L ±1.75 mm, D/V -4.7 mm. The pipette was held in place for an additional 4 min. Two weeks after viral injection, diphtheria toxin (DT, 100 μg/kg, Sigma D0564) was intraperitoneally injected to ablate cholinergic neurons expressing DTR. Two weeks later, behavioral tests were performed.

### Chemogenetic modulation of cholinergic neuron activity

In order to test the role of cholinergic neuron activity during coordinated motor performance, the designer receptor exclusively activated by designer drug (DREADD) system was used. Twenty hemizygous *ChAT-Cre* mice were bilaterally injected with Cre-dependent inhibitory DREADD fused with mCherry reporter AAV8-hsyn-DIO-hM4Di-mCherry (1×10^13^ VG/ml; Addgene, 44362) or control virus (AAV8-hsyn-DIO-mCherry) (1×10^13^ VG/ml; Addgene, 50459) into 3 NBM/SI sites (350 nl/site) as with AAV-DTR. After 3 weeks to allow for viral expression, the DREADD ligand JHU37160 (J60, Hello Bio) was administered via intraperitoneal injection at a dose of 1.0 mg/kg 30 minutes prior to rotarod training.

### Accelerating rotarod test

Mice were habituated on the rotarod (Ugo Basile, 47650) at a speed of 4 rpm for 60 s before testing. For each trial, the rotarod accelerated from 5 to 60 rpm over 300 s. In SAP and DT experiments, mice were trained for 3 trials/day over 3 days with a 30 min interval between trials. For DREADD experiments, mice were tested 30 min after IP injection of DREADD ligand, 3 trials/day over 3 days without an interval between trials. In alternate rotarod training, mice were habituated at 4 rpm for 60 s on a different rotarod (Med Associates, Inc. ENV-577M), with trials accelerating from 4 to 40 rpm over 5 min. The latency to fall after the onset of acceleration during each trial was recorded for each mouse. Individual trials were stopped, and the duration was recorded if mice could not run with consecutive rotations or failed to stay on the rotarod.

### Recessed forelimb reach task

Mice were food restricted to 80–90% of their free-feeding bodyweight before training. An acrylic behavior box with three slots (7 mm wide) on the left, middle, and right sides of the front wall was used to train the mice. A recessed hole (3 mm wide, 2 mm deep) at 12 mm from the inside wall of the box was used to hold a 20 mg flavored food pellet (Bioserv, F05301). The dominant forelimb for reaching was identified during one session of pellet reaching. Once the dominant forelimb was determined, it was trained over a total of 14 daily sessions consisting of 25 trials each. Mice often become sated and less likely to perform skilled reach after 25 trials. A trial was counted as a success if the mouse successfully grasped, retrieved, and ate the pellet. Only trials with pellet contact were counted. The success rate was defined as the percentage of trials with successful retrieval and eating.

### Nestlet shredding

1.2 g nestlets were placed into each test cage and the mass of remaining intact nestlet was measured at several time points over a one-week period.

### Open field test

Mice were placed in a chamber (L×W×H: 30 cm×22.5 cm×25 cm) and allowed to explore for 5 min. Behavior was recorded from the top of the chamber at 48 fps (GoPro, HERO3+) and analyzed by MatLab software (Autotyping15.04 source code) (Patel et al., 2014). Thigmotaxis was defined as the percentage of time that mice spent within two inches of the arena walls.

### Wire hanging test

Mice were individually placed on a wire mesh grid. Once the animal grasped the grid with all four paws and appeared stable, the mesh was inverted and placed atop an open chamber. The duration that the mice were able to hang from the grid was recorded. A soft blanket was placed at the bottom of the chamber to avoid animal injury. Each animal was tested 3 times and the longest duration was used as the animal’s latency to fall.

### Retrograde NBM/SI labeling

Retrograde tracing was performed in 8-week-old C57BL/6J mice (*n* = 4) in order to label basal forebrain neurons projecting to specific cortical areas. Alexa Fluor 488, 555, or 647 conjugated Cholera Toxin B subunit (CTB) (1% wt/vol, Molecular Probes) was bilaterally injected to mPFC (A/P +1.9 mm, M/L ±0.3 mm, D/V -1.5 mm), motor cortex (+0.5 mm, ±1.5 mm, -0.6 mm), or visual cortex (−3.3 mm, ±2.3 mm, -0.6 mm), respectively. A burr hole was drilled over each corresponding area and 100 nl tracer was injected at each site at a rate of 40 nl/min. To prevent backflow, the pipette was left in the brain for 5 minutes after injection. Mice were sacrificed 1 week later. Transverse brain sections were cut coronally at 40μm of thickness. ChAT immunostaining was performed to label cholinergic neurons. Some sections were counterstained with DAPI. Sections containing injection sites were imaged at 10x on a Leica SP8 confocal microscopy with parameters adjusted based on the intensity of expression and background fluorescence (tile scan). Images containing cholinergic neurons were acquired using 10x objective.

### Histology

Mice were anesthetized with ketamine/xylazine cocktail [150 mg/kg; 15 mg/kg] and transcardially perfused with ice-cold PBS followed by 4% paraformaldehyde (PFA). Brains were harvested and post-fixed in 4% PFA overnight at 4°C, cryoprotected by immersion in 30% sucrose in 0.1M PBS for 2 days, and transversely cryosectioned (40 μm thick) using a Leica cryostat. Free floating sections were permeabilized with 0.3% Triton X-100 in PBS for 30 min at room temperature. After blocking with 10% donkey serum, sections were incubated with primary antibodies for 2 days at 4°C. Primary antibodies used: goat anti-ChAT (1:100, Millipore, AB144P), rabbit anti-p75 (1:100, Advanced Targeting Systems, AB-N01AP), rabbit anti-GFP (1:1500, Invitrogen, A6455), rabbit anti-DsRed antibodies (1:3000, Takara, 632496). Sections were washed three times in PBS, followed by incubation with fluorescently conjugated secondary antibodies (1:200, Jackson ImmunoResearch) for 1.5 hours at room temperature. Nuclei were labeled with a 10-minute incubation with DAPI [1 μg/ml] in PBS. Images were acquired on a Leica SP8 confocal microscope with 10x or 20x objectives under constant imaging parameters. Total CTB-labeled NBM/SI neurons in serial sections were manually counted.

### Statistics

Rotarod, skilled pellet reach, and nestlet shredding behavior tests were analyzed using two-way repeated measures analysis of variance (ANOVA) with post-hoc Sidak’s comparison test using GraphPad Prism 9.0. Two-tailed t-tests were used to compare differences between two groups. Two-way ANOVA with post hoc Dunnett’s multiple comparisons test was used to compare cholinergic fibers among different groups. All behavior and analysis were performed in a double-blinded manner. For all figures, \**P* < 0.05; \*\**P* < 0.01; \*\*\**P* < 0.001.

## Results

### Ablation of basal forebrain cholinergic neurons impairs coordinated motor training

In order to ablate basal forebrain cholinergic neurons, we injected C57BL/6J mice bilaterally with anti-p75 conjugated saporin (p75-SAP) into the nucleus basalis of Meynert (NBM) and substantia innominata (SI); control mice were injected with IgG-conjugated saporin (IgG-SAP) (Figure 1A). p75-SAP targets cholinergic neurons in the basal forebrain as they selectively express the low-affinity nerve growth factor p75NTR receptor. p75-SAP infusion resulted in nearly complete loss of NBM/SI cholinergic neurons (Figure 1B,C; *n* = 8, two-tailed t-test, *P* < 0.001). Motor coordination was tested using an accelerating rotarod. Mice were trained over 9 trials starting at 5 rpm with a constant acceleration over 5 minutes to 60 rpm. Both male and female mice injected with p75-SAP exhibited worse performance on accelerating rotarod compared to IgG injected controls, with significantly shorter latency to fall (Figure 1D,E; *n* = 8, two-way repeated measures ANOVA, *P* = 0.004). These results confirmed the findings from a pilot study in which mice trained under alternate conditions (4-40 rpm over 5 min) on a different rotarod showed impaired performance following depletion of basal forebrain cholinergic neurons by p75-SAP injection, relative to IgG-SAP controls (Figure 1—figure supplement 1; *n* = 6, two-way ANOVA, *P* < 0.001). In addition to rotarod deficits, animals with cholinergic neuron ablation showed a reduction in repetitive, nestlet-shredding behavior. Control mice quickly tear up nestlet bedding material placed in their cages while p75-SAP injected mice showed dramatically reduced nesting behavior as determined by measuring the mass of nestlets over 7 days (Figure 1F,G; two-way ANOVA, *P* < 0.003).

**Figure 1.**
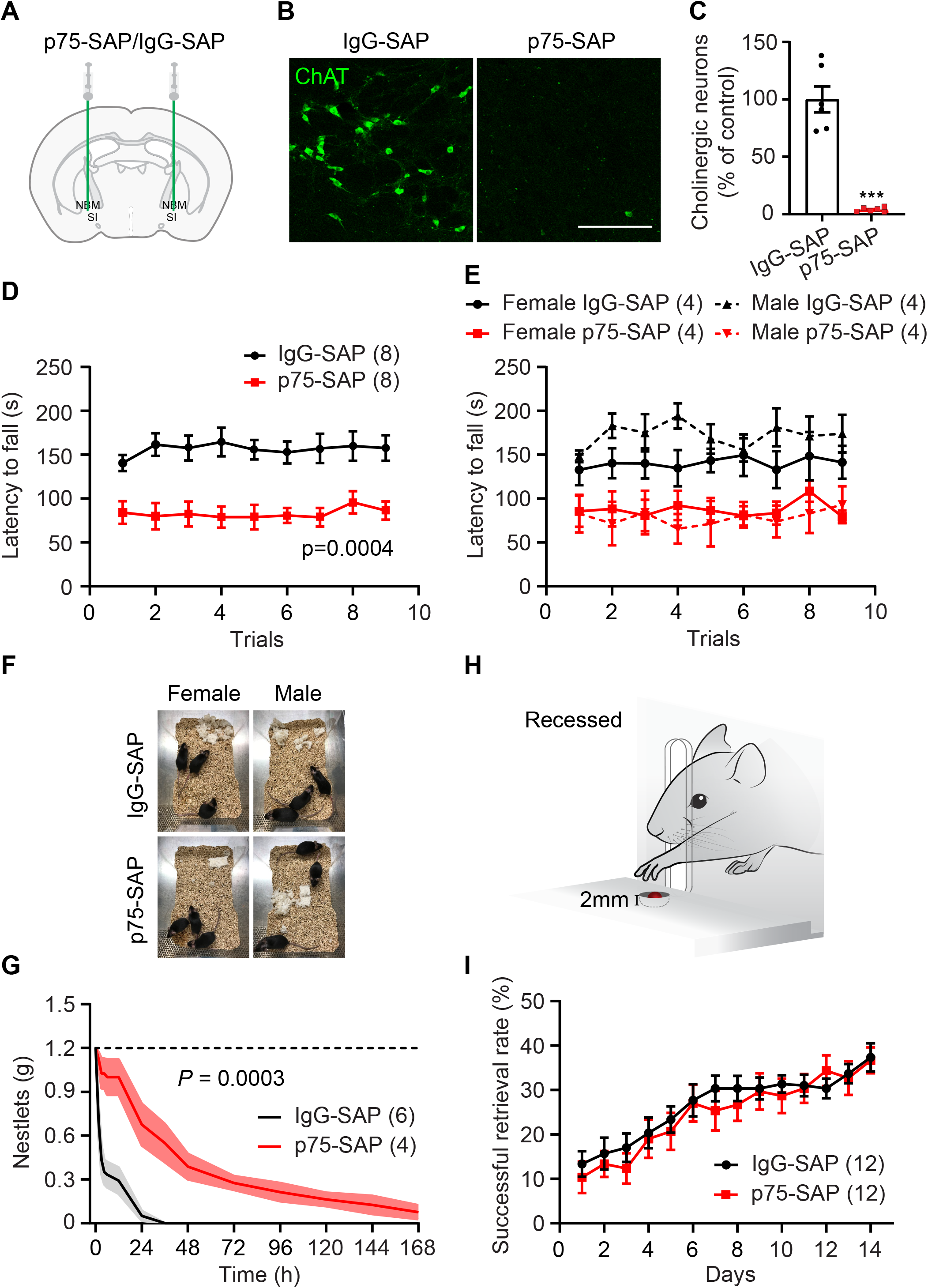
Pharmacological ablation of basal forebrain cholinergic neurons impairs coordinated motor learning. (A) NBM/SI basal forebrain cholinergic neuron targeting with anti-p75 conjugated saporin or control IgG-saporin. (B) NBM/SI cholinergic neurons following IgG-SAP or p75-SAP injection (scale bar = 200 μm). (C) Quantification of ChAT immunostained neurons in NBM/SI basal forebrain in animals with IgG-SAP (control) or p75-SAP injection (two-tailed t-test, \*\*\**P* < 0.001). (D) Ablation of cholinergic neurons by p75-SAP severely impaired rotarod training performance (repeated measures ANOVA, *P* = 0.0004, F (1, 14) = 21.31). (E) p75-SAP effects were independent of sex. (F) Nestlet shredding behavior in female and male mice injected with IgG-SAP (control) or p75-SAP. (G) Quantification of remaining, intact nestlets over time (*n* is number of cages, repeated measures ANOVA, *P* = 0.0003, F (1, 8) = 35.79). (H) Illustration of the recessed forelimb reach task. (I) Learning of the recessed forelimb reach task was unimpaired following p75-SAP injection (repeated measures ANOVA, *P* > 0.05, F (1, 22) = 0.2983). Data presented as mean ± s.e.m., *n* in parentheses is number of mice unless otherwise indicated.

Mice were also trained to perform a recessed version of the single pellet reach task over 2 weeks. The single pellet reach task is frequently used in rat models to study motor learning (Bova et al., 2019; Chen et al., 2014; Zemmar et al., 2015); however, many mice fail to show an improvement on the standard pellet reach task, exhibiting an essentially flat learning curve across training (Chen et al., 2014). We therefore used a modified, recessed version of the skilled reach task in which mice have to retrieve a food pellet from a concave depression (Figure 1H). On the standard task, the initial success rate was 27 ± 5%, while after two weeks of training this had only increased to 37 ± 4% (Figure 1—figure supplement 2A,B). The resulting increase was not statistically significant (paired t-test, *P* = 0.15). Separate mice trained on the recessed forelimb reach task had a lower initial success rate at 17 ± 3%, but a more consistent improvement, increasing to 41 ± 3% with training (paired t-test, *P* < 0.0001). 43% of mice trained on the standard forelimb reach task failed to improve by more than 15% over the course of two weeks (Figure 1—figure supplement 2C), while 93% of mice trained on the recessed forelimb reach task demonstrated improvements with training. This recessed version of the single pellet reach task allowed us to assess skilled motor learning in our study. We found that p75-SAP injection did not impair the learning of the recessed forelimb reach task, as both groups exhibited a similar improvement in performance over the course of training (Figure 1I).

We tested the overall health of mice after cholinergic ablation. At two weeks after saporin injection, mice injected with p75-SAP exhibited a slight but significant decrease in the body mass (Figure 1—figure supplement 3A; *n* = 8, paired t-test, *P* = 0.002). Overall animal strength was unaffected as determined using a wire hanging test. Both groups showed a similar time to fall, regardless of gender (Figure 1—figure supplement 3B,C). General activity was tested in an open field (Figure 1—figure supplement 3D). p75-SAP injected mice exhibited a reduced total walking distance compared to IgG injected controls (Figure 1—figure supplement 3E, two-tailed t-test, *P* =0.042). As mice navigated the open field, they largely remained close to the walls (thigmotaxis), indicative of a normal level of anxiety. No difference in thigmotaxis was observed between groups (Figure 1—figure supplement 3F).

### Genetic lesion of basal forebrain cholinergic neurons impairs coordinated motor training

We next used a targeted genetic approach to deplete basal forebrain cholinergic neurons. *ChAT-Cre::Ai14* mice were bilaterally injected into NBM/SI with AAV encoding the diphtheria toxin receptor in a Cre-dependent manner (AAV/DJ8-FLEX-DTR-EYFP). Control mice were injected with AAV expressing only Cre-dependent EYFP fluorescent reporter (Figure 2A). Diphtheria toxin (DT) injection ablated nearly all cholinergic neurons in the NBM and SI areas in mice transduced with AAV-FLEX-DTR-EYFP (Figure 2B,C and Figure 2—figure supplement 1A). Motor cortex and mPFC received strong cholinergic innervation from NBM/SI areas, as exhibited in mice injected with control virus (Figure 2D). Two weeks after DT injection, AAV-DIO-DTR-EYFP mice showed a mild, but significant decrease in body mass (Figure 2—figure supplement 1B; *n* = 8, paired t-test, *P* < 0.001). DT-injected control animals showed no reduction in body mass. Similar to the effects of p75-SAP NBM/SI lesion, DT injection into AAV-DIO-DTR-EYFP transduced *ChAT-Cre::Ai14* mice resulted in severely impaired performance on the accelerating rotarod (5-60 rpm over 5 min) (Figure 2 E,F; *n* = 8, ANOVA, *P* = 0.001). Furthermore, *ChAT-Cre::Ai14* mice transduced with DTR virus exhibited dramatically reduced nestlet shredding behavior after DT treatment (Figure 2G; ANOVA, *P* < 0.001). DT injection had no effect on the learning of the recessed forelimb reach task (Figure 2—figure supplement 1C-E).

**Figure 2.**
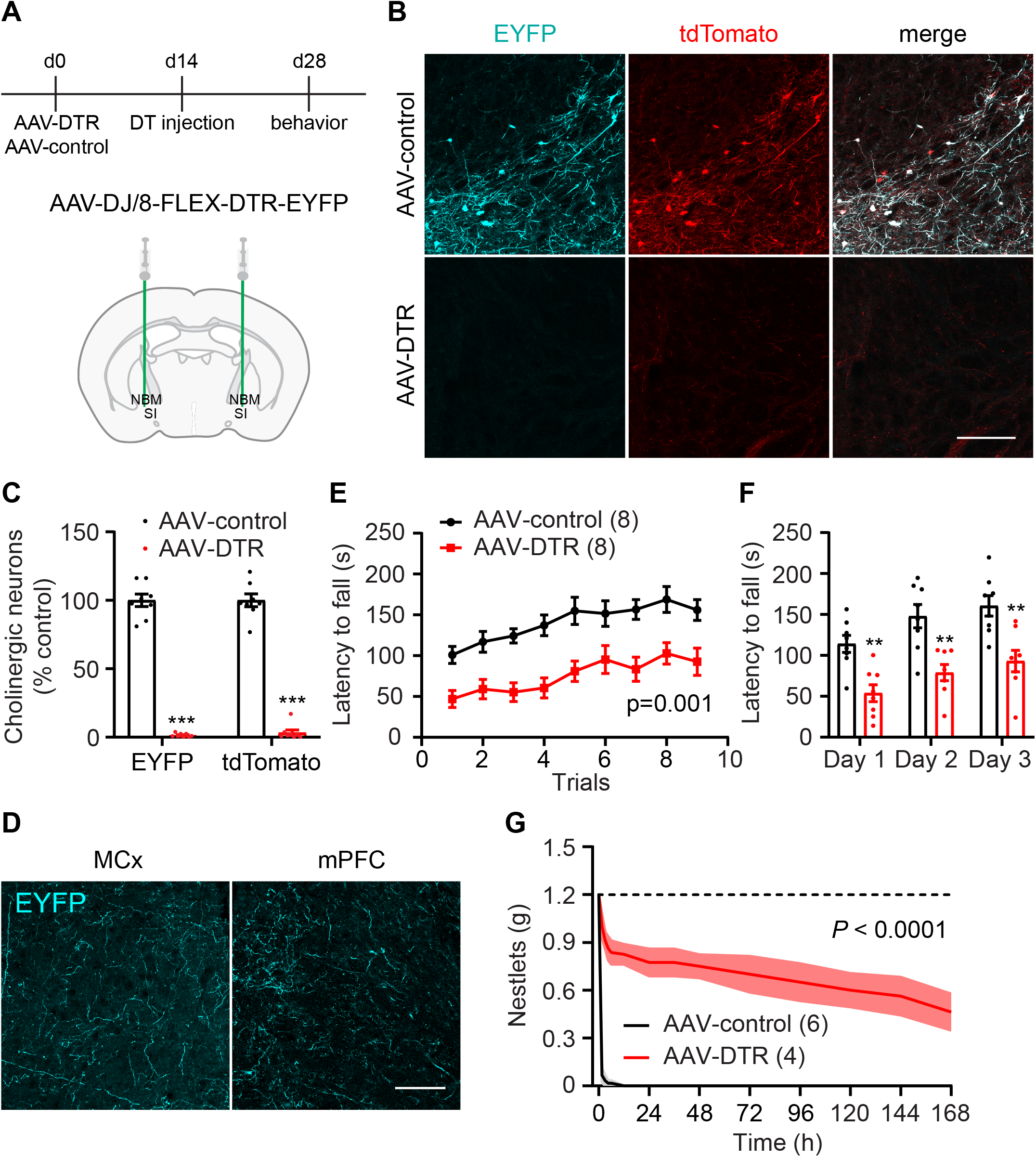
Genetic depletion of cholinergic neurons impairs coordinated motor learning. (A) Timeline outlining genetic ablation of cholinergic neurons in hemizygous *ChAT-Cre::Ai14* mice followed by behavior tests (top panel). NBM/SI basal forebrain cholinergic neuron transduction with AAV-DTR or control EYFP. (B) NBM/SI cholinergic neurons expressing tdTomato in mice injected with AAV-FLEX-EYFP (control) or AAV-FLEX-DTR-EYFP (scale bar = 200 μm). (C) Quantification of EYFP or tdTomato positive neurons in NBM/SI basal forebrain (two-tailed t-test, \*\*\**P* < 0.001). (D) EYFP-labeled cholinergic fibers in motor cortex and mPFC after AAV-FLEX-EYFP transduction of cholinergic neurons in *ChAT-Cre* mice (scale bar = 100 μm). (E) Genetic ablation of cholinergic neurons in basal forebrain severely impaired performance on rotarod training (repeated-measures ANOVA, *P* = 0.001, F (1, 14) = 17.30). (F) Rotarod latencies averaged for 3 trials each day (repeated measures ANOVA with post-hoc Sidak’s comparison test, \*\**P* < 0.01). (G) Quantification of remaining, intact nestlets over time (*n* is number of cages, repeated measures ANOVA, *P* < 0.0001, F (1, 8) = 117.4). Data presented as mean ± s.e.m., *n* in parentheses is number of mice unless otherwise indicated.

### Chemogenetic silencing of cholinergic neurons impairs coordinated motor training

In order to test the role of cholinergic neuron activity during the acquisition of coordinated motor behavior, the designer receptor exclusively activated by designer drug (DREADD) system was used. *ChAT-Cre* mice were bilaterally injected into NBM/SI with Cre-dependent inhibitory DREADD AAV-DIO-hM4Di-mCherry or control virus (AAV-DIO-mCherry) (Figure 3A). After 3 weeks to allow for viral expression (Figure 3B), rotarod training was performed 30 min after intraperitoneal injection of the DREADD ligand JHU37160 (J60, 1 mg/kg). A total of 9 trials were performed on the accelerating rotarod (5-60 rpm over 5 min). Inactivation of cholinergic neurons during behavior significantly impaired rotarod performance (Figure 3C,D; *n* = 10, ANOVA, *P* = 0.0033).

**Figure 3.**
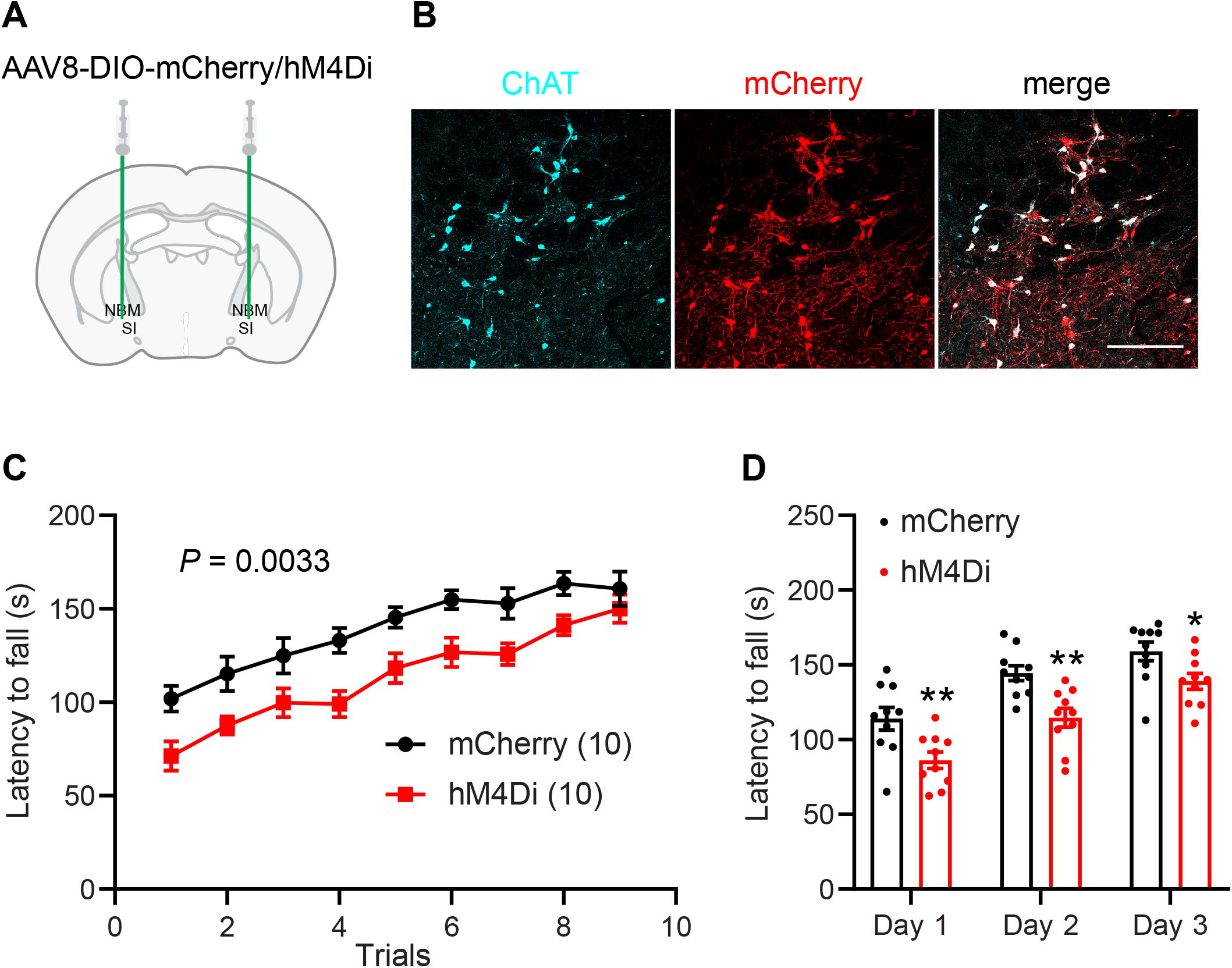
Chemogenetic silencing of cholinergic neurons impairs coordinated motor performance. (A) NBM/SI basal forebrain cholinergic neuron transduction with AAV-DIO-hM4Di-mCherry in hemizygous *ChAT-Cre* mice. (B) hM4Di-mCherry fusion protein expression in ChAT positive neurons (scale bar = 200 μm). (C) JHU37160 delivered 30 minutes prior to behavior severely impaired performance on rotarod training in mice expressing hM4Di in NBM/SI cholinergic neurons (repeated-measures ANOVA, *P* = 0.0033, F (1, 18) = 11.44). (D) Rotarod latencies averaged for 3 trials each day (repeated measures ANOVA with post hoc Sidak’s comparison test, \**P* < 0.05, \*\**P* < 0.01). Data presented as mean ± s.e.m..

### Cholinergic projections to both motor cortex and mPFC are required for coordinated motor training

As basal forebrain cholinergic neurons project extensively throughout the cortex, we selectively targeted cholinergic inputs to distinct cortical areas in order to determine the target locus of acetylcholine action during coordinated motor learning. We injected p75-SAP directly into motor areas required for coordinated motor learning, mPFC and motor cortex, as well as into visual cortex, which is not anticipated to be involved in the task (Figure 4 A,B and Figure 4—figure supplement 1). Control animals were injected with IgG-SAP into mPFC, motor, or visual cortices. IgG-SAP injection into different targets had no effect on rotarod behavior and data from these animals were pooled. Two weeks after SAP injections, mice were trained on the accelerating rotarod. Depletion of cholinergic input to both mPFC and motor cortex produced significant deficits in rotarod performance, compared to control IgG-SAP injected animals (Figure 4C and Figure 4—figure supplement 2A; ANOVA, *P* = 0.0011). Visual cortex injection of p75-SAP had mild but not significant effects on behavior. Unlike global depletion of NBM/SI cholinergic neurons, there was no obvious effect of local ablation on nestlet shredding behavior (Figure 4—figure supplement 2B). Mice with targeted ablation of cholinergic neurons were then trained on skilled behavior. Similar to global ablation of NBM/SI cholinergic neurons, target-specific ablation of cholinergic neurons had no effect on training of the recessed forelimb reach task (Figure 4—figure supplement 2C). These data indicate that cholinergic basal forebrain neurons regulate motor coordination through modulation of both primary and associative motor circuits (Figure 4D).

**Figure 4.**
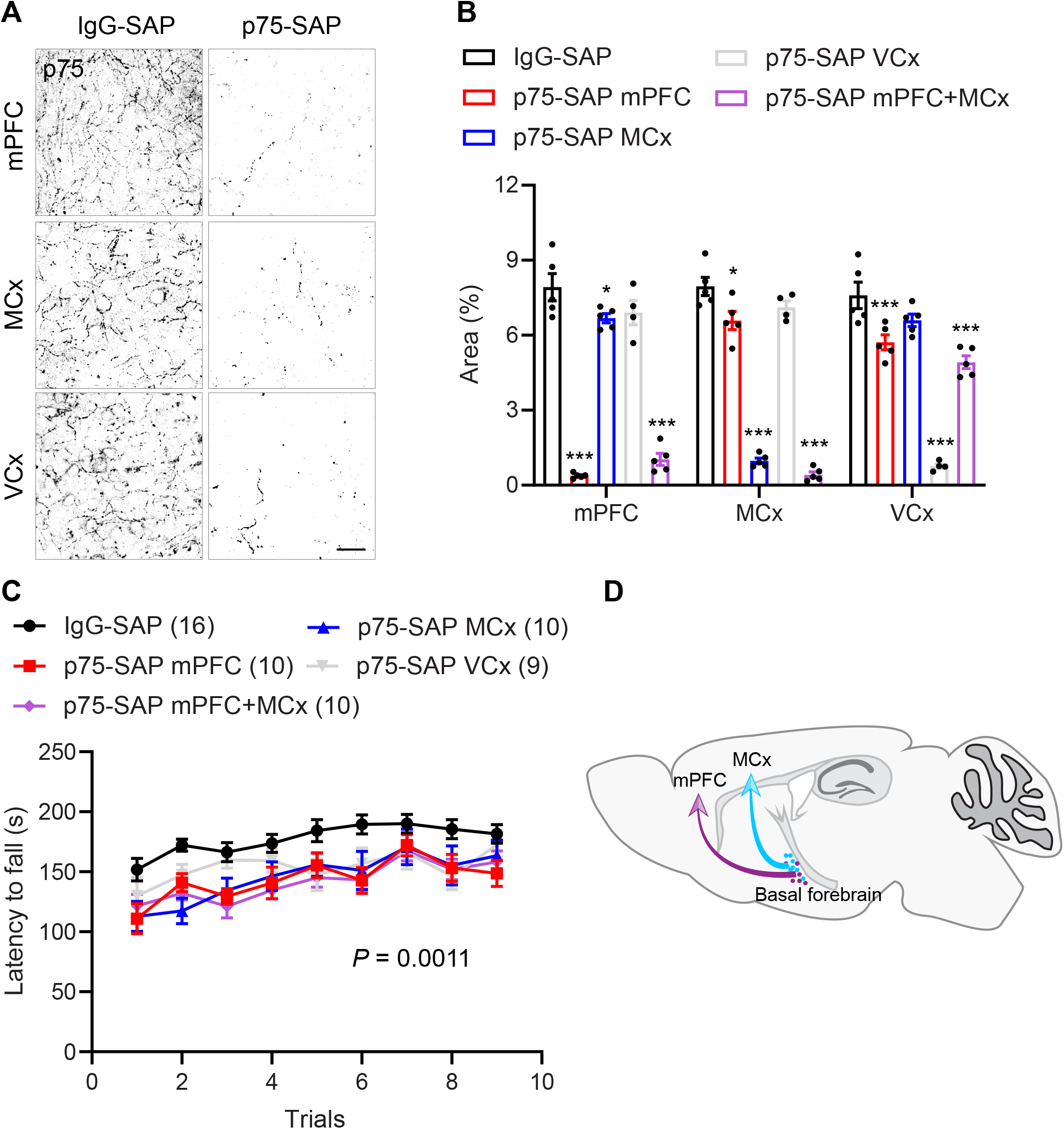
Cholinergic inputs to both mPFC and motor cortex are required for coordinated motor training. (A) Cholinergic innervation in mPFC, motor cortex (MCx), and visual cortex (VCx) after selective targeting with IgG-SAP or p75-SAP (scale bar = 20 μm). (B) Quantification of p75 positive cholinergic fibers in mPFC, MCx, and VCx in response to selective ablation of cholinergic inputs (two-way ANOVA with post hoc Dunnett’s multiple comparisons test, \**P* < 0.05, \*\*\**P* < 0.01). (C) Depletion of cholinergic inputs to either mPFC, MCx, or both mPFC and MCx significantly impaired performance on rotarod behavior training. Depletion of cholinergic input to visual cortex (VCx) had a partial impact on rotarod performance (repeated-measures ANOVA, *P* = 0.0011, F (4, 50) = 5.382.). (D) Schematic illustrating cholinergic innervation of distant motor centers involved in coordinated motor learning. Data presented as mean ± s.e.m..

In order to determine if collateral branches of NBM/SI neurons innervate the anatomically isolated regions tested above, we injected unique fluorescently conjugated retrograde tracer CTB into mPFC (488 nm), motor cortex (555 nm), and visual cortex (647 nm, Figure 5A,B). Overall, 58% of labeled neurons in basal forebrain were ChAT-positive. The majority of CTB-labeled basal forebrain input to motor and visual cortices were from cholinergic neurons at 71.2±6.1 and 57.2±2.0%, respectively. In mPFC, the percentage of cholinergic neurons was 49.4±1.4% (Figure 5C). Retrogradely labeled, non-cholinergic, basal forebrain neurons include GABAergic or glutamatergic neurons (Gritti et al., 2003; Henny and Jones, 2008; Kim et al., 2015). The majority of cholinergic neurons (84.4±2.0%) were labeled with a single CTB tracer, indicating that there was no overlap in their axonal projections within the labeled regions (Figure 5 D,E). Although the sites of retrograde tracer injection were distal to each other, we did observe overlapping labeling of a small population of basal forebrain cholinergic neurons. Double-labeled neurons projecting to mPFC and motor cortex represented 4.1±1.4% of NBM/SI cholinergic neurons, motor and visual cortices 3.6±1.4%, and mPFC and visual cortex 5.9±2.4%. Cholinergic neurons labeled by three colors of CTB were observed rarely (2.1±1.5%). The shared innervation may, at least in part, explain the mild effects of cholinergic input to visual cortex on rotarod training.

**Figure 5.**
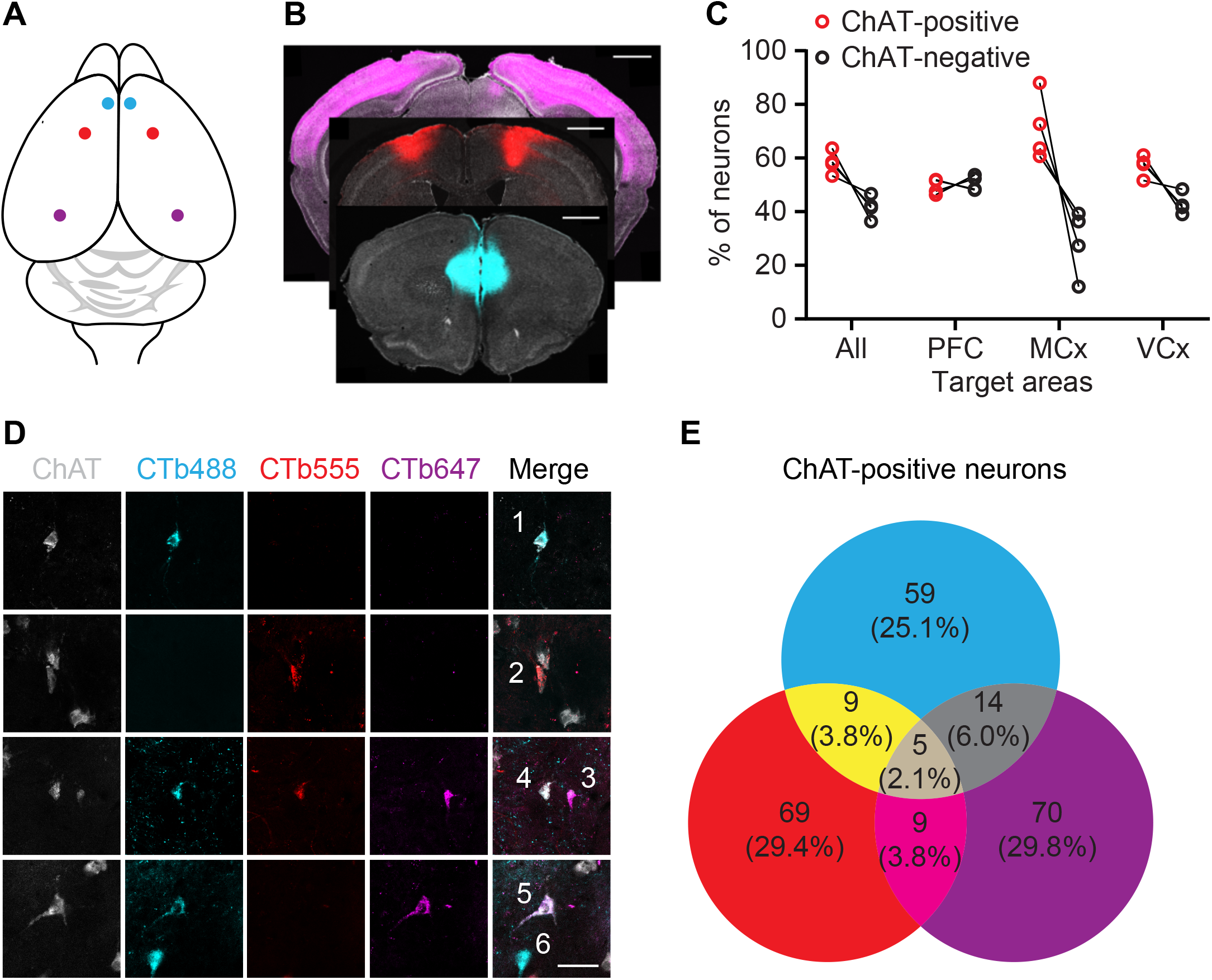
Retrograde labeling of NBM/SI cholinergic projections to cortex. (A) Illustration of retrograde tracing strategy: CTB conjugated to Alexa Fluor 488, 555, or 647 was bilaterally injected to mPFC (cyan), motor cortex (red), or visual cortex (purple), respectively. (B) CTB injection sites: mPFC (cyan), motor cortex (red), and visual cortex (purple). Sections were stained with DAPI (gray) (scale bar = 1 mm). (C) Quantification of cholinergic and non-cholinergic neurons projecting to cerebral cortices. (D) Retrograde labeling of NBM/SI with CTB conjugates. Neurons 1, 2 and 3 projected to one of these three cortices. Neuron 4 projected to mPFC and motor cortex. Neuron 5 projected to mPFC and visual cortex. Neuron 6 is non-cholinergic (scale bar = 40 μm). (E) Quantification of total CTB-labeled cholinergic NBM/SI neurons.

### Basal forebrain cholinergic neurons are not required for execution of previously trained coordinated motor behavior

To determine whether basal forebrain cholinergic neurons are simply required for the execution rotarod behavior, rather than for the learning of the coordinated motor task, we trained intact mice prior to NBM/SI cholinergic neuron ablation. Mice were first trained on the accelerating rotarod for 9 trials and then randomly assigned to treatment with either p75-SAP or control IgG-SAP injection into NBM/SI. Two weeks later, the retention of trained rotarod behavior was tested (Figure 6A). Execution of the previously trained rotarod behavior was unaffected in both groups, with comparable fall latencies before and after SAP injection (Figure 6B,C). Next, we used a genetic approach to test the effects of cholinergic ablation on coordinated motor task retention. *ChAT-Cre* transgenic mice were injected with AAV-DIO-DTR-EYFP or control virus. Ten days later, mice were trained through 9 trials on the accelerating rotarod prior to intraperitoneal injection of DT to ablate cholinergic neurons (Figure 6D). DTR and EYFP control expressing mice exhibited similar rotarod performance both before and after DT injection (Figure 6E,F). These results demonstrate that basal forebrain cholinergic signaling is not needed for the execution of previously learned coordinated motor behavior.

**Figure 6.**
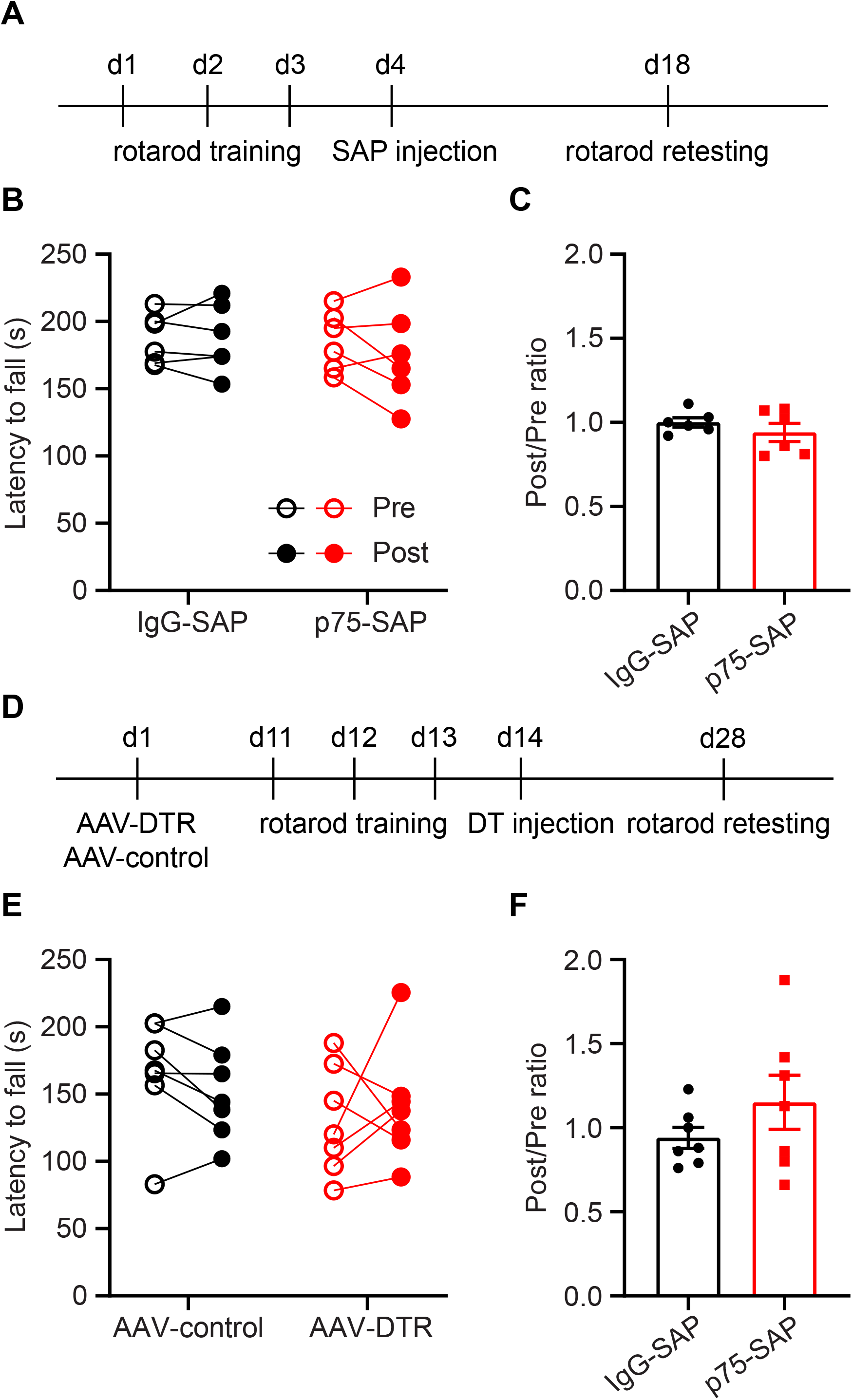
Basal forebrain cholinergic neurons are not required for execution of previously learned coordinated motor behavior. (A) Timeline outlining experimental details of cholinergic ablation by p75-SAP following rotarod training. Retention testing was performed 2 weeks after saporin injection. (B) Pharmacological ablation of cholinergic neurons did not affect the performance on the previously learned rotarod behavior (paired t-tests, *P* > 0.05 for both groups). (C) The ratio of latencies to fall before and after cholinergic ablation (Post/Pre) (two-tailed t-test, *P* = 0.34). (D) Timeline outlining experimental details of cholinergic ablation by AAV-DTR following rotarod training. Rotarod retention testing was performed 2 weeks after DT injection. (E) Genetic ablation of cholinergic neurons did not affect the performance on the previously learned rotarod behavior (paired t-tests, *P* > 0.05 for both groups). (F) The ratio of latencies to fall before and after DT-induced cholinergic ablation (Post/Pre) (two-tailed t-test, p = 0.24). Data presented as mean ± s.e.m..

## Discussion

Our findings in this study demonstrate that basal forebrain cholinergic neurons exert control over coordinated motor learning. Ablation or inhibition of these cholinergic neurons impairs coordinated task acquisition but does not affect execution of previously learned coordinated behavior. These findings are important as the release of acetylcholine rapidly modulates neuronal excitability, circuit dynamics, and cortical coding; all processes required for processing complex sensory information, cognition, and attention (Ballinger et al., 2016; Goard and Dan, 2009; Metherate et al., 1992; Villano et al., 2017). Baso-cortical cholinergic pathways are critical neuromodulators of auditory, visual, olfactory, and somatosensory perception. In primary auditory cortex, pairing electrical stimulation of NBM with a specific auditory stimulus leads to circuit remodeling (Bakin and Weinberger, 1996; Kilgard and Merzenich, 1998). Within S1, increased cholinergic signaling suppresses slow spontaneous activity and drives the depolarization of layer 2/3 neurons during whisking (Eggermann et al., 2014). Modulating the activity of basal forebrain cholinergic neurons alters visual and olfactory discrimination capabilities (Goard and Dan, 2009; Ma and Luo, 2012; Pinto et al., 2013). In addition to sensory coding, the cholinergic input to primary visual cortex is also required for the acquisition, but not the expression, of reward timing in the primary visual cortex (Chubykin et al., 2013; Liu et al., 2015). Cholinergic neurons also disinhibit layer 1 interneurons in auditory cortex and mediate associative fear learning (Letzkus et al., 2011). Our study raises the possibility that baso-cortical cholinergic input rapidly modulates the active state of motor networks during coordinated motor learning.

In addition to mPFC and sensorimotor cortex, learned motor movements rely on the coordinated output of thalamus, basal ganglia, cerebellum, and brainstem motor centers. Training drives a shift in the contributions of cortical and subcortical motor circuits, with an early instructional role for motor cortex supporting the development of independent movement initiation by the basal ganglia (Kawai et al., 2015). Indeed, the contribution of the motor cortex to learned movements appears to decline over time with continued training (Hwang et al., 2019). We demonstrate clearly that basal forebrain cholinergic signaling is critical for the early instructional phase of coordinated motor learning. As we found that cholinergic inputs to both mPFC and motor cortex are required for coordinated motor acquisition, it is likely that cholinergic neuromodulation plays a critical role in the coactivation of mPFC-DMS and M1-DLS circuits during early stage rotarod learning (Kupferschmidt et al., 2017). As training progresses and instructional input from cortical projections is no longer driving experience-dependent refinement of striatal motor circuits (Wolff et al., 2019), cholinergic neuromodulation of cortical motor centers is no longer required. This is supported by our findings that ablation of cholinergic neurons after rotarod training elicited no effect on performance of the previously trained task.

Basal forebrain cholinergic input to M1 has been shown to be a key modulator of the cortical plasticity mechanisms that underlie skilled forelimb reach training in rats (Conner et al., 2003). Several lines of evidence implicate cholinergic neuromodulation in the large-scale cortical changes that occur following injury or motor learning in other species (Conner et al., 2005; Juliano et al., 1991). While we did not assess cortical reorganization in our studies, our data showed that skilled motor learning on a recessed version of the single pellet reach task was unaffected following cholinergic ablation in mice. These results may owe to species differences in the refinement of cortical mechanisms, or perhaps to differences in the acetylcholine-dependent development of cortical motor representations (Ramanathan et al., 2015). In fact, mice often fail to demonstrate the learning curves exhibited by rats in the standard single pellet reach task used in earlier studies. Typically a large percentage of mice will exhibit a high proficiency on the task on the first days of training and show no subsequent improvement, or even a decline in performance over time (Figure 1—figure supplement 2) (Chen et al., 2014). The modified, recessed forelimb reach task appears to be more difficult for mice, and reliably results in performance improvements with training. Another critical difference observed was that our studies showed a role for mPFC cholinergic inputs in coordinated motor task acquisition, while p75-SAP targeting of mPFC had no effect on forelimb reach training in rats (Conner et al., 2010). This is consistent with the coordination of activity across mPFC and M1 that others have shown in rotarod training (Kupferschmidt et al., 2017). Whether there are specific basal forebrain circuits, or other neuromodulatory mechanisms, in mice that differentially regulate the acquisition of coordinated and skilled tasks is a critical question that requires further study.

Beyond the effects on cortical plasticity, acetylcholine signaling plays a key role in cognitive processes by mediating attention (Sarter and Lustig, 2019; Thiele and Bellgrove, 2018). Within the visual cortex, attention drives increased firing rates via acetylcholine signaling through muscarinic receptors in non-human primates (Herrero et al., 2008). Within humans, numerous studies have suggested a critical role for external focus of attention in skilled motor learning (Lewthwaite and Wulf, 2017; Lohse et al., 2014; Song, 2019; Wulf, 2013). External focus of attention is tightly linked to practice variability and task improvements that occur through movement revision during motor learning (Chua et al., 2019). Meanwhile, divided attention impairs task learning by reducing movement adaptations across trials (Song, 2019). The effects of external focus of attention are robust, driving motor learning both in neurotypical individuals as well as in individuals with intellectual disabilities (Chiviacowsky et al., 2013). The central role of attention in the acquisition of motor skills supports the idea that cognitive components are indispensable in the adjustment of motor output and training-induced improvements in motor performance (Gallivan et al., 2018; McDougle et al., 2016).

As we observed a critical role for cholinergic input to mPFC in coordinated motor learning, it may be that disruption of circuit level mediators of attention impair motor learning. Prefrontal cortex in primates mediates top-down attentional control over sensory systems through direct input to inferotemporal cortex (Baluch and Itti, 2011; Miller and Buschman, 2013; Petrides, 2000). Within mPFC, attention drives increases in synchronous activity of fast-spiking parvalbumin inhibitory interneurons and increased gamma oscillations (Kim et al., 2016). Acetylcholine signaling through muscarinic receptors in mPFC is required for cue-mediated increases in gamma oscillations (Howe et al., 2017; Parikh et al., 2007). On a cued discrimination task, optogenetic activation of basal forebrain cholinergic neurons has been shown to enhance sensitivity to short duration cues, but also to increase incorrect attempts in the absence of cues (Gritton et al., 2016). In contrast, silencing of cholinergic inputs impairs cue detection (Gritton et al., 2016). A role for cholinergic modulation of attention during coordinated motor learning may, in part, explain motor deficits in neurological disorders otherwise characterized by cognitive dysfunction. While the hallmarks of neurodegenerative disorders such as Alzheimer’s disease (AD) are cognitive impairments, the neuropathological effects on attention in AD (Miloyan et al., 2013) may explain pre-clinical motor decline in AD individuals (Albers et al., 2015) and coordinated motor learning impairments in AD mouse models (O’Leary et al., 2018). Our results may provide insight into the relevance of motor and sensory dysfunction and pre-clinical AD prognosis.

## Acknowledgements

We thank Amanda Bernstein for assisting in behavioral tests and data analysis, Anne Marchildon for animal breeding and genotyping, and Sydney Agger for illustrations. We also thank Rachel Garn for early contributions to the manuscript. We acknowledge IDDRC Neuroconnectivity Core at Baylor College of Medicine and the viral core at Boston Children’s Hospital for producing DTR virus. pAAV-hSyn-DIO-hM4Di-mCherry was generated by Bryan Roth (Addgene plasmid # 44362). This work was supported by Burke Foundation and the National Institutes of Health Common Fund DP2 NS106663 to EH, New York State Department of Health SCIRB postdoctoral fellowship (C32633GG) to YL.

## Author contributions

YL and EH designed the study; YL performed the experiments. YL and EH wrote the manuscript.

## Competing interests

The authors declare that no competing interests exist.

## Figure Legends

**Figure 1—figure supplement 1.**
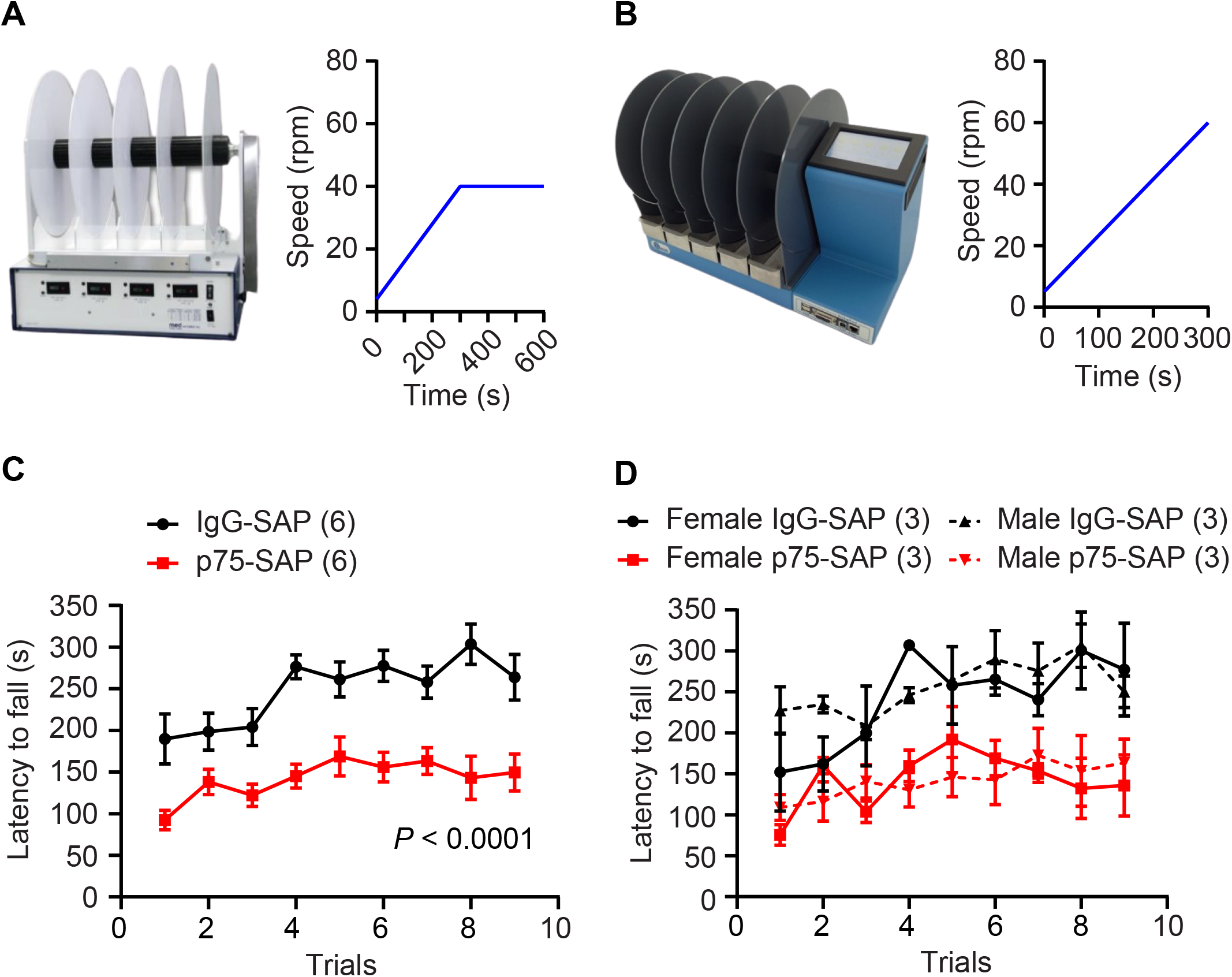
Accelerating rotarod tests. (A) In an alternate rotarod training paradigm, C57BL/6J mice were trained on a Med Associates, Inc. ENV-577M rotarod with trials accelerating from 4 to 40 rpm over 5 min followed by an additional 5 min at 40 rpm. (B) Ugo Basile 47650 rotarod used in main figures; the training paradigm was continuous acceleration from 5 to 60 rpm over 5 min. (C) Ablation of cholinergic neurons by p75-SAP severely impaired alternate rotarod training performance (repeated-measures ANOVA, *P* < 0.0001, F (1, 10) = 42.74). (D) p75-SAP effects were independent of sex. Data presented as mean ± s.e.m..

**Figure 1—figure supplement 2.**
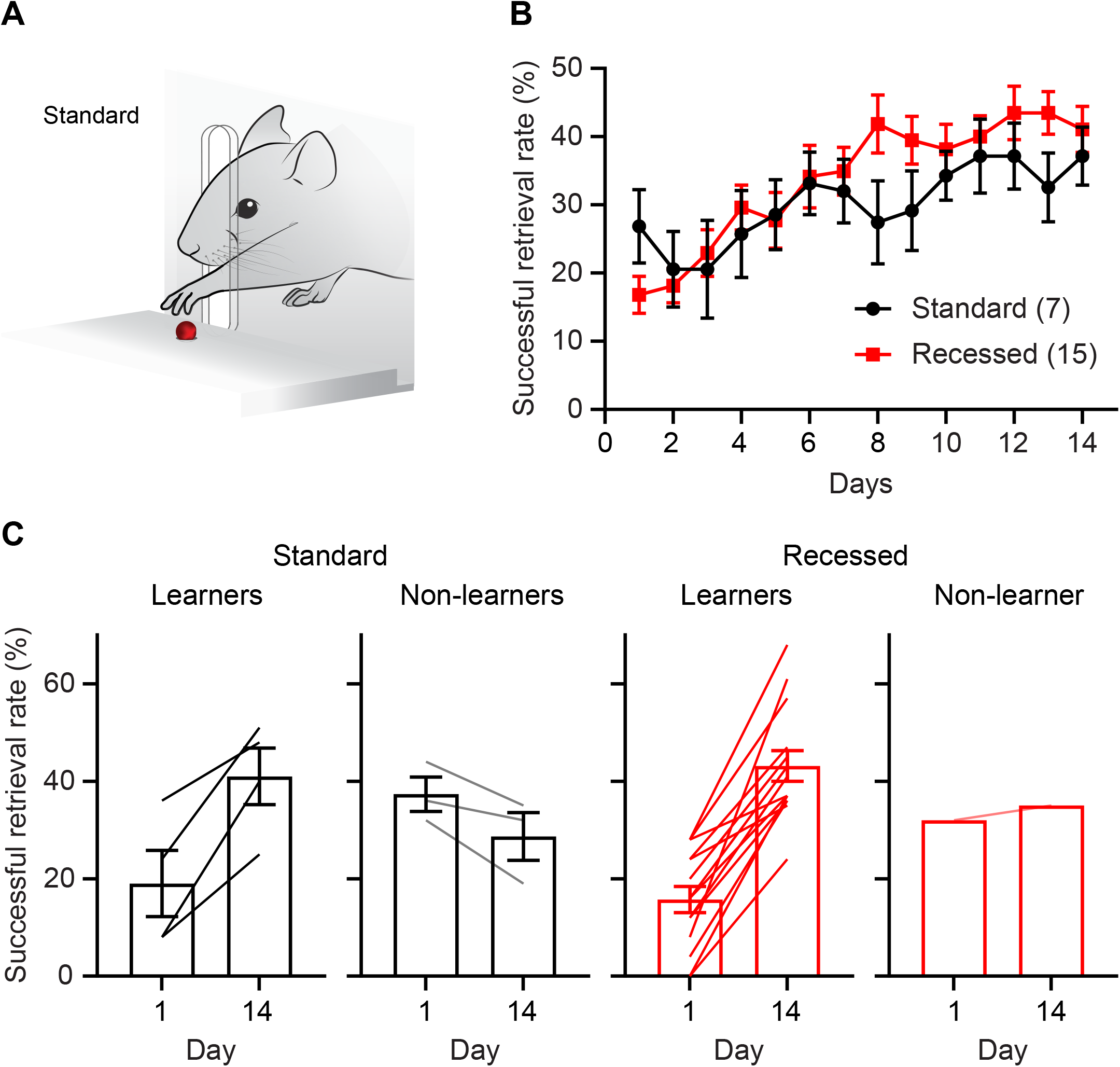
Mice exhibit more consistent learning on the recessed forelimb reach task. (A) Illustration depicting the standard skilled pellet reach behavior. (B) Successful retrieval rate significantly improves over 2 weeks of behavioral training on the recessed version compared to the standard task. (C) The majority of mice exhibit learning over the course of training on the recessed version of the task (14/15), in contrast to the standard task (4/7). Data presented as mean ± s.e.m..

**Figure 1—figure supplement 3.**
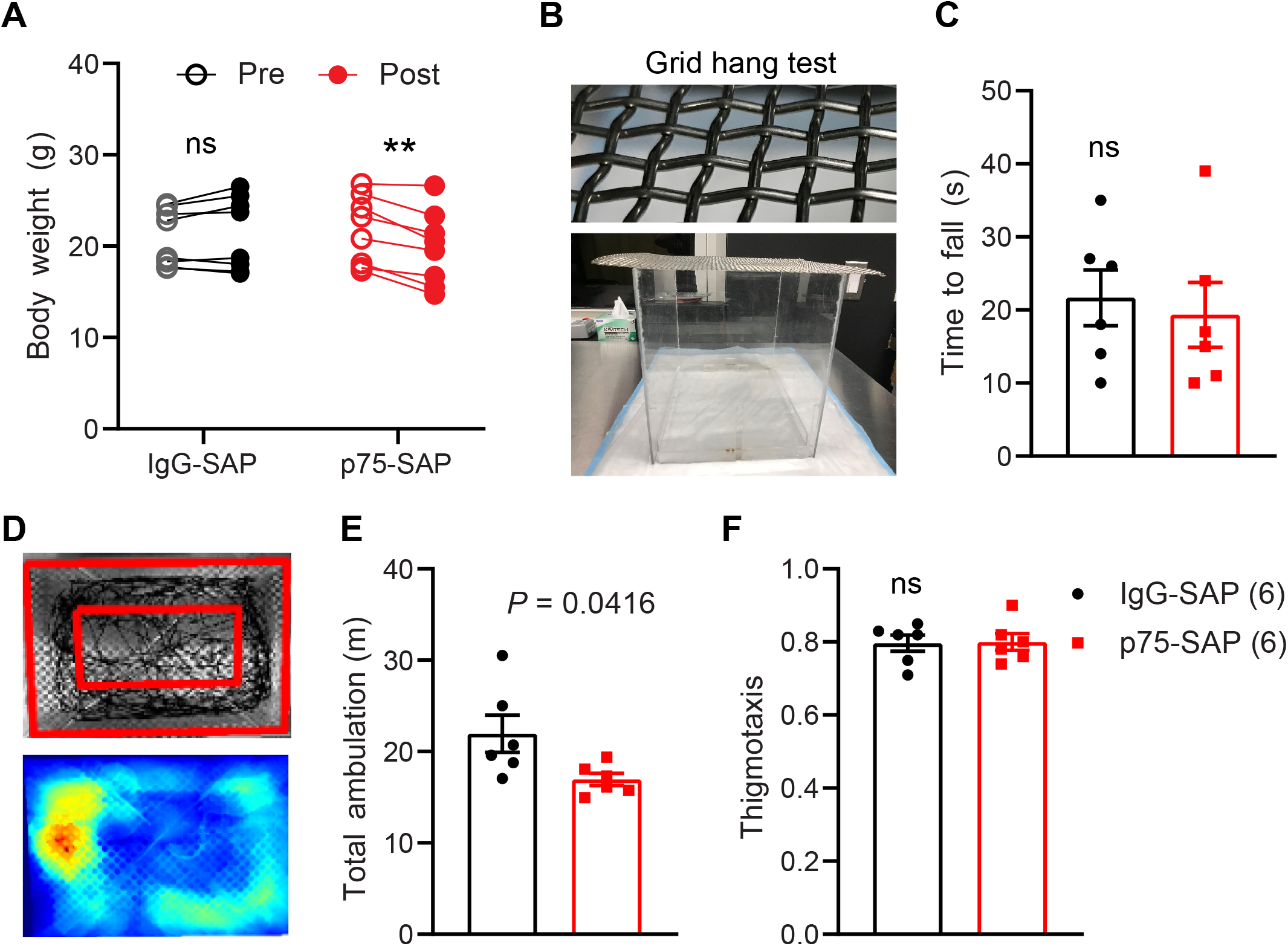
General effects of cholinergic ablation by p75-SAP. (A) Mice injected with p75-SAP had a slight but significant decrease in body mass (paired t-test, **p < 0.01). (B) Grid hanging test. (C) Animal strength was similar between treatment groups (two-tailed t-test, *P* = 0.70). (D) Representative sample of the open field test showing walking trace of one mouse (top panel); outer red rectangle marked the boundary of walking space. The distance between 2 rectangles is 2 inches. Heat map of walking trace (bottom panel). (E) Total walking distance was shorter in mice injected with p75-SAP (two-tailed t-test, *P* = 0.042). (F) Thigmotaxis was similar between treatment groups Data presented as mean ± s.e.m..

**Figure 2—figure supplement 1.**
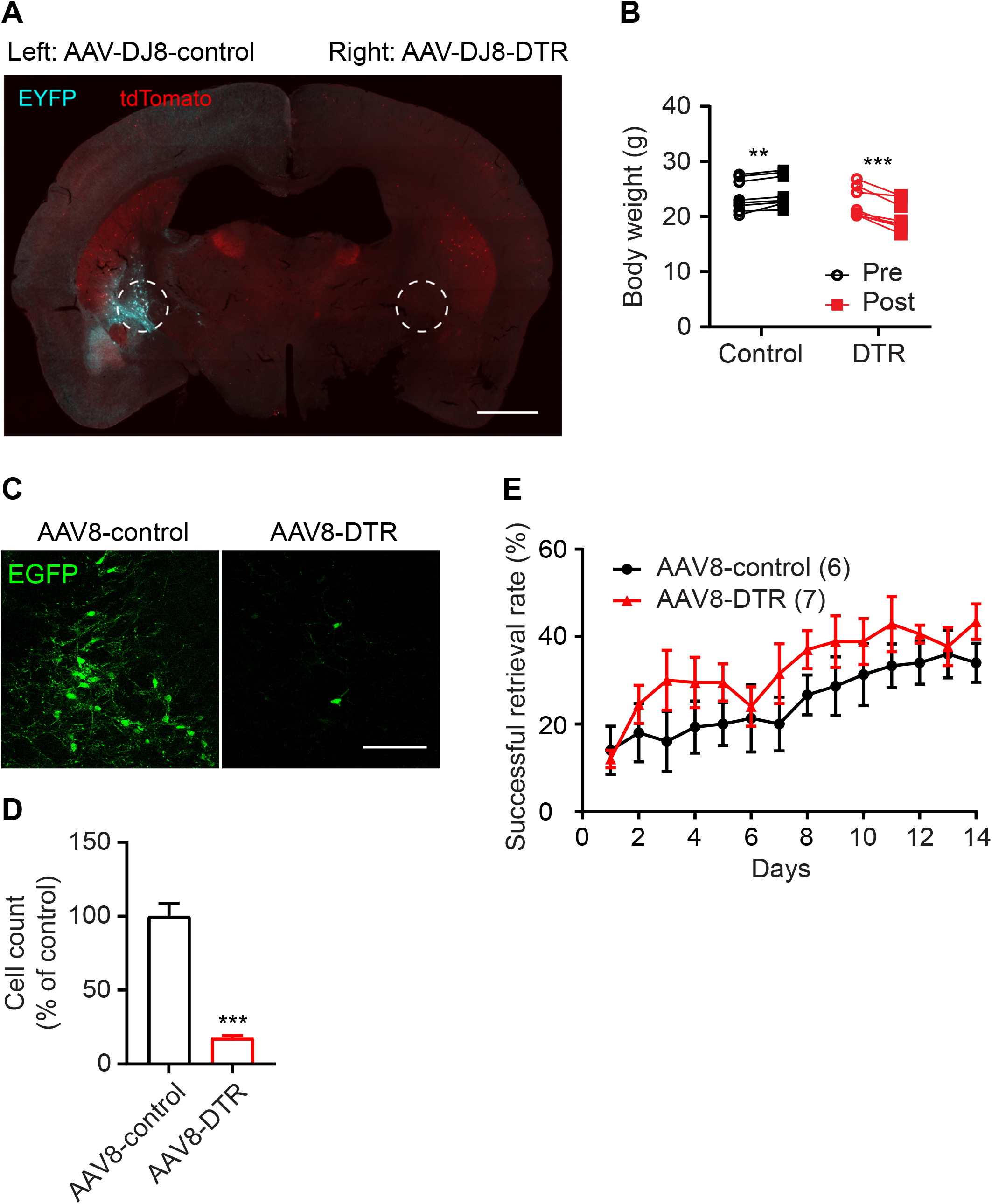
DT-induced ablation of DTR-expressing cholinergic neurons. (A) Transverse section from a *ChAT-Cre::Ai14* mouse transduced with control AAV-FLEX-EYFP (left) or AAV-FLEX-DTR-EYFP (right) showing targeted depletion of NBM/SI cholinergic neurons at 2 weeks after DT injection (scale bar = 1 mm). (B) DTR expression in NBM/SI cholinergic neurons leads to a slight but significant decrease in the body mass at 2 weeks after DT injection (paired t-tests, ** *P* <0.01, *** *P* < 0.001). (C-E) In a subset of experiments, hemizygous *ChAT-Cre* mice were co-injected with AAV8-syn-FLEX-DTR and AAV2-syn-FLEX-EGFP. Two weeks later, mice were injected with DT. Confocal images showed EGFP expression in mice with or without transduction of DTR virus (C). Scale = 200 μm. (D) Quantification of the number of EGFP positive neurons in basal forebrain of animals injected with AAV8-DTR as compared to control animals (unpaired t-test, *** *P* < 0.001). (E) Genetic ablation of cholinergic neurons did not impair learning of recessed forelimb reach task (repeated-measures ANOVA, *P* = 0.1429, F (1, 11) = 2.49). Data presented as mean ± SEM.

**Figure 4—figure supplement 1.**
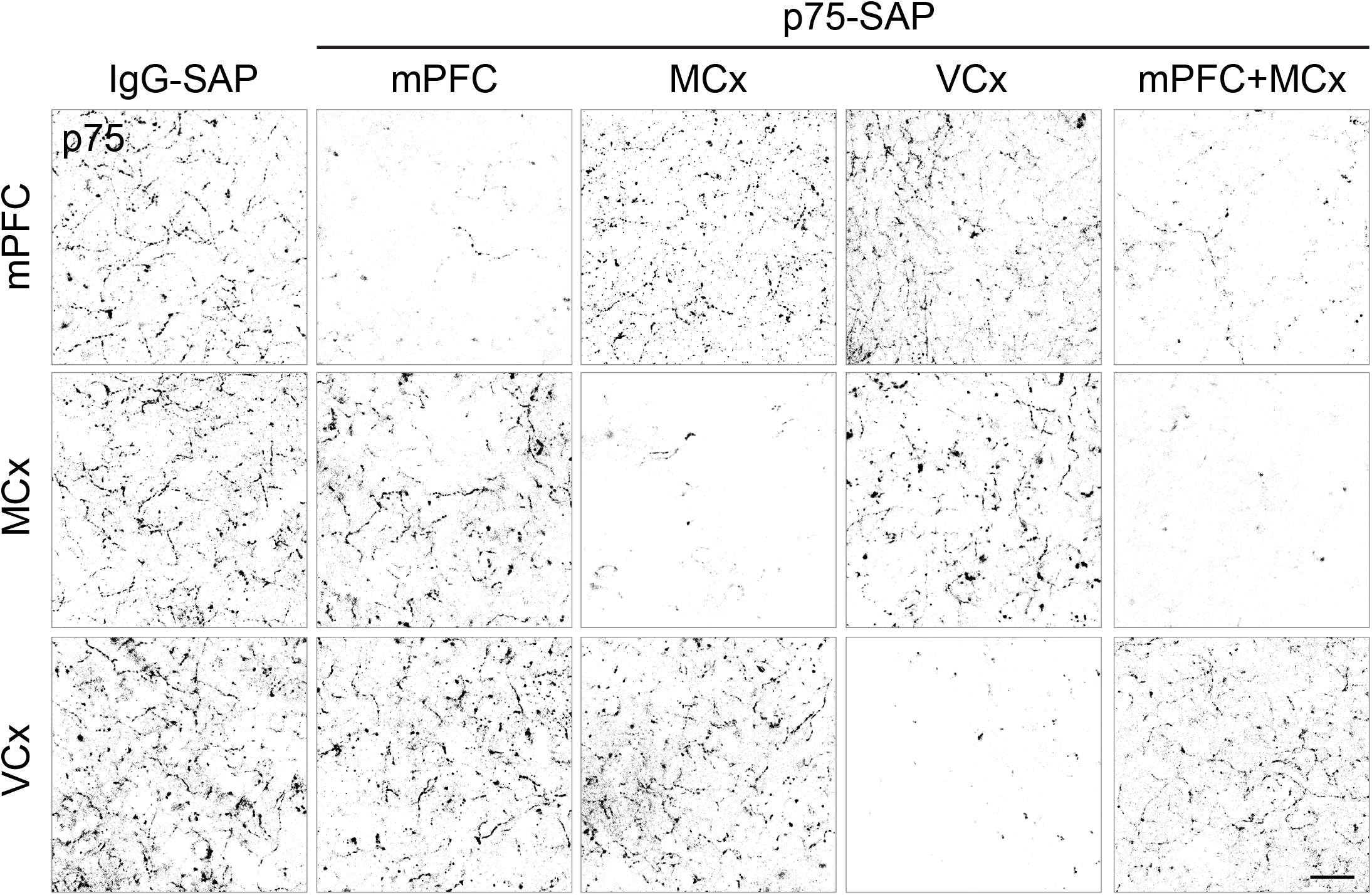
Depletion of cholinergic innervation in distinct cortical areas. p75 immunostaining of cholinergic fibers in mPFC, MCx, and VCx from mice with local injection of p75-SAP or control IgG-SAP (scale bar = 20 μm).

**Figure 4—figure supplement 2.**
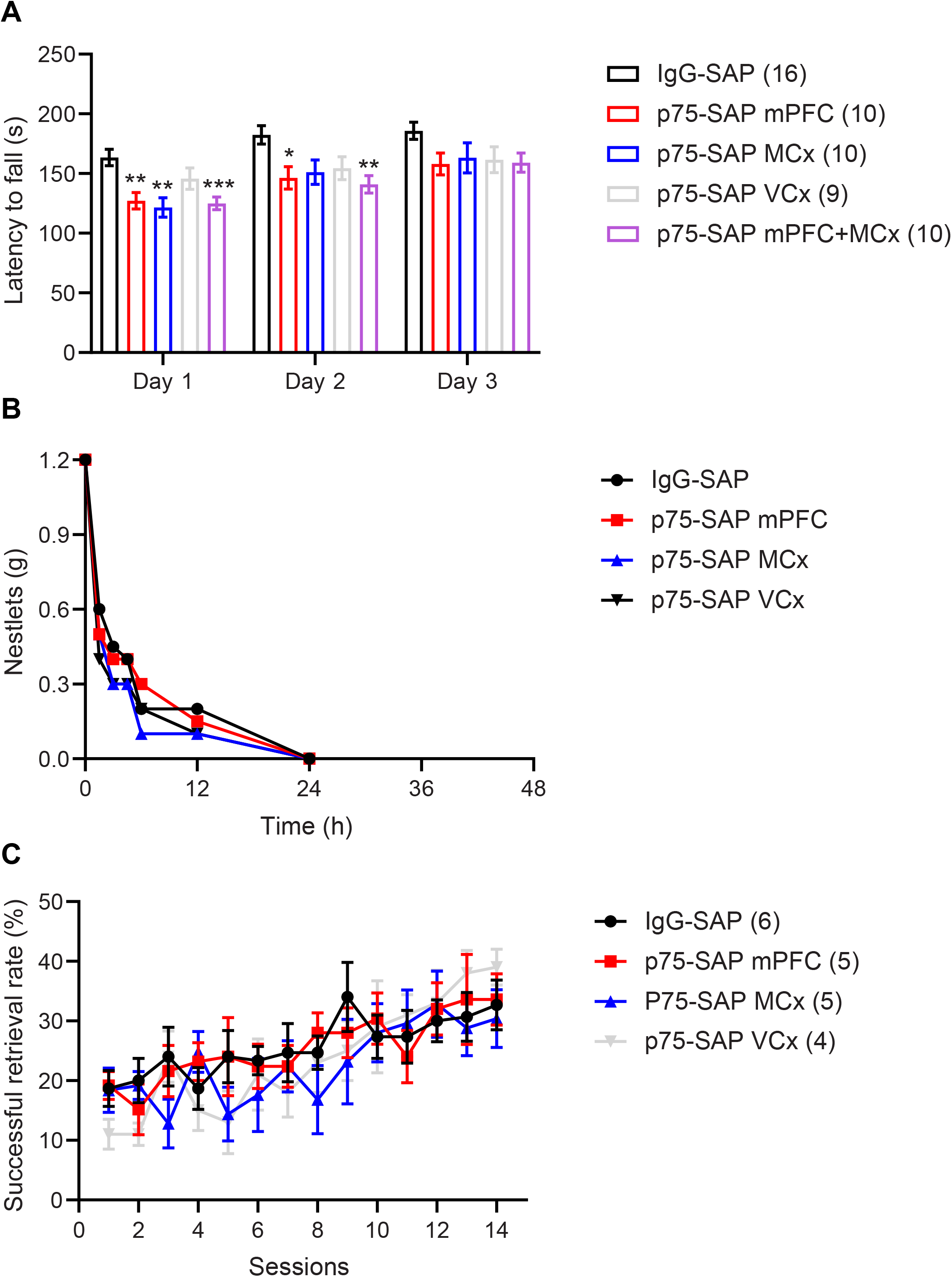
General effects of depletion of cholinergic inputs to specific cortical areas. (A) Rotarod latencies averaged for 3 trials each day (repeated measures ANOVA with post hoc Sidak’s comparison test. \**P* < 0.05, \*\**P* < 0.01, \*\*\**P* < 0.001). (B,C) Depletion of cholinergic inputs to mPFC, motor cortex (MCx), or visual cortex (VCx) has no effect on nestlet shredding behavior (B) or recessed forelimb reach task (C).

## Notes

### Competing Interest Statement

The authors have declared no competing interest.

